# Topology-Driven Discovery of Transmembrane Protein *S*-Palmitoylation

**DOI:** 10.1101/2024.09.08.611865

**Authors:** Michael T. Forrester, Jacob R. Egol, Sinan Ozbay, Rohit Singh, Purushothama Rao Tata

## Abstract

Protein *S*-palmitoylation is a reversible lipophilic posttranslational modification regulating a diverse number of signaling pathways. Within transmembrane proteins (TMPs), *S*-palmitoylation is implicated in conditions from inflammatory disorders to respiratory viral infections. Many small-scale experiments have observed *S*-palmitoylation at juxtamembrane Cys residues. However, most large-scale *S*-palmitoyl discovery efforts rely on trypsin-based proteomics within which hydrophobic juxtamembrane regions are likely underrepresented. Machine learning– by virtue of its freedom from experimental constraints – is particularly well suited to address this discovery gap surrounding TMP *S*-palmitoylation. Utilizing a UniProt-derived feature set, a gradient boosted machine learning tool (TopoPalmTree) was constructed and applied to a holdout dataset of viral *S*-palmitoylated proteins. Upon application to the mouse TMP proteome, 1591 putative *S*-palmitoyl sites (i.e. not listed in SwissPalm or UniProt) were identified. Two lung-expressed *S*-palmitoyl candidates (synaptobrevin Vamp5 and water channel Aquaporin-5) were experimentally assessed. Finally, TopoPalmTree was used for rational design of an *S*-palmitoyl site on KDEL-Receptor 2. This readily interpretable model aligns the innumerable small-scale experiments observing juxtamembrane *S*-palmitoylation into a proteomic tool for TMP *S*-palmitoyl discovery and design, thus facilitating future investigations of this important modification.

## INTRODUCTION

Protein *S*-palmitoylation is a reversible posttranslational modification that allows for hydrophobic “tuning” of specific protein regions. For membrane-associated proteins such as the Ras family of GTPases, *S*-palmitoylation facilitates proper membrane localization and signaling(1). Within transmembrane proteins (TMPs), *S*-palmitoylation is believed to modulate protein conformation relative to the lipid bilayer(2), ultimately contributing to diverse biological outputs from extracellular signal transduction to endocytic sorting. In the case of pathogenic coronaviruses, *S*-palmitoylation of the viral Spike protein (a TMP) is essential for proper envelope geometry and viral replication(3, 4).

Gradient boosted trees (GBTs) are a type of supervised iterative machine learning that employs gradient descent to improve the performance of successive trees(5). These flexible algorithms can handle different types of data, be tuned to minimize overfitting and maintain a structure amenable to model interpretation. Within molecular biology, gradient boosted machines have been employed for predictive models of *O*-phosphorylation(6, 7), protein-small molecule(8) and -DNA(9) interactions. Gradient boosted trees, in particular, frequently outperform other methods such as Naïve Bayes, support vector machines and logistic regression(10) as manifested by the widespread success of GBTs in machine learning competitions.

Despite the biological relevance of TMP *S*-palmitoylation, our knowledge about molecular determinants of *S*-palmitoylation has primarily relied on small-scale studies focused on individual proteins. Early experiments suggested a narrow range of distance from the lipid bilayer in dictating *S*-palmitoylation of an individual Cys residue on ER protein p63, whereas adjacent amino acids were less critical(11). A globally applicable motif for TMP *S*-palmitoylation has not been found(12) and typical molecular cues for *S*-palmitoylation, such as *N*-myristoylation or C-terminal isoprenylation, do not appear operative on TMPs. Despite the enigmatic nature of TMP *S*-palmitoylation, this modification has been shown to dictate localization of TMPs to specific membrane microdomains(13, 14), receptor internalization from the plasma membrane(15, 16), proper tilting of transmembrane domains relative to the lipid bilayer(17, 18) and assembly into organized protein complexes(2, 12).

Gradient boosted trees are known to excel with complex mixtures of categorical and continuous features, yet GBTs remain to be utilized for *S*-palmitoyl site discovery(19). Further, most established *S*-palmitoyl inference tools primarily rely on sequence information(20–22) rather than the topological features supported by small-scale experiments. Motivated by statistical(23) and experimental(24, 25) investigations suggesting unique physiochemical determinants for TMP *S*-palmitoylation, we have sidestepped the traditional one-size-fits-all model and constructed a TMP-specific algorithm for *S*-palmitoyl discovery.

## RESULTS

### Juxtamembrane regions are under-represented in large scale *S*-palmitoyl data

Twenty to thirty percent of mammalian open reading frames encode transmembrane proteins (TMPs), many of which are signaling receptors and clinically relevant pharmacological targets. These proteins exhibit regions of elevated hydrophobicity that are underrepresented in bottom-up proteomics due to their relative resistance to trypsinization and poor aqueous solubility(26, 27). Given the propensity of Cys residues to undergo *S*-palmitoylation in these hydrophobic juxtamembrane regions, we hypothesized that *S*-palmitoyl datasets derived primarily from bottom-up proteomics might encompass only a small fraction of *bona fide S*-palmitoylated sites.

To evaluate this question, the mouse transmembrane proteome was subjected to *in silico* trypsinization followed by biophysical peptide analysis. Of 63380 Cys-containing peptides, 4357 match to reported *S*-acylated sites in SwissPalm with 955 of these sites being located on a TMP. The physical properties of all Cys-containing peptides were plotted with respect to molecular weight and mean hydrophobicity (based on Kyte-Doolittle values). As shown in Figure 1A/B, the majority (80.3%) of TMP *S*-palmitoyl sites in SwissPalm fall into the ideal range of detectability defined as MW 700 – 3000 Da and mean hydrophobicity of −2 to +1. However, categorizing the same 63380 Cys residues by membrane proximity – defined as being within 20 amino acids of a transmembrane domain – reveals that only 22.5% of membrane-proximal Cys-containing peptides fall within this ideal range (Figure 1C/D). These findings suggest that membrane-proximal *S*-palmitoyl “hot spots” may be underrepresented in high throughput experiments, the vast majority of which employ trypsin-based bottom-up proteomics. While experimentally tailored proteolysis can improve TMP recall in bottom-up proteomics, *in silico* approaches can completely circumvent these peptide-level biophysical constraints.

**Figure 1.**
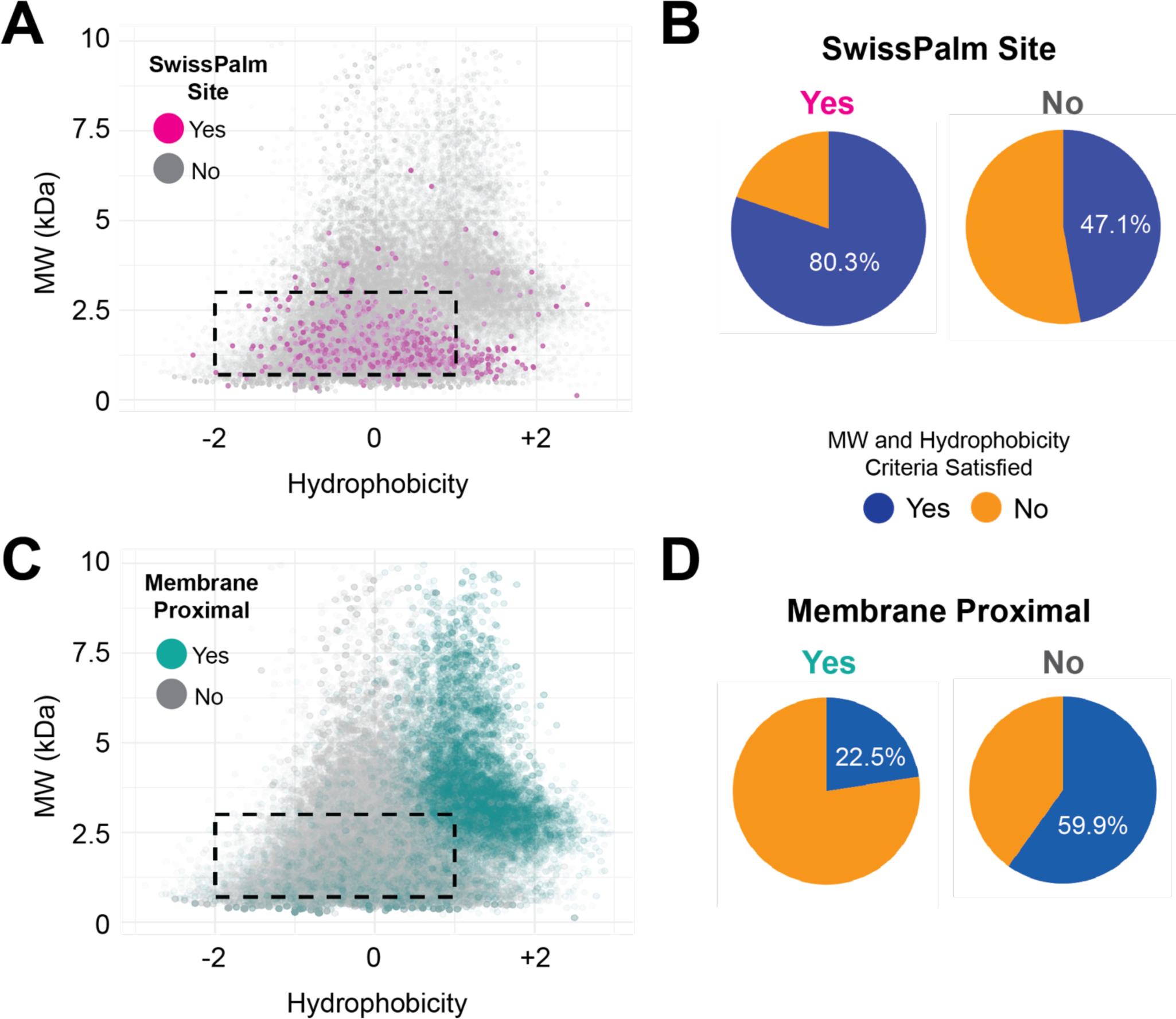
Experimental “detectability” of TMP *S*-palmitoyl sites. The murine transmembrane proteome was subjected to *in silico* trypsinization, followed by labeling of Cys-containing peptides based on whether they are (A & B) reported as *S*-palmitoylated in SwissPalm (magenta) or (C & D) within 20 amino acids of a transmembrane domain and thus proximal to the lipid bilayer (teal). The dashed rectangle represents a general detectible region for bottom-up proteomics of 700 – 3000 Da and mean Kyte-Doolittle hydrophobicity of −2 to +1. The pie charts indicate the fraction of Cys-containing peptides that fall within the detectable range based on whether the Cys sites are (B) reported in SwissPalm or (D) juxtamembrane in location.

### Establishment of a high fidelity *S*-palmitoyl TMP training dataset

To isolate a training dataset with reliable and relatively complete representation of TMP *S*-palmitoyl sites, large scale proteomics data was set aside in favor of data primarily obtained by two rigorous methods: site-directed mutagenesis and ^3^H-palmitate radiolabeling. Additional data (4.9% of the positive class) was obtained from native LC-MS(24) and considered “positive” for every Cys site at least 25% *S*-palmitoylated. After filtering and collecting available topological data from UniProt, 446 sites of *S*-palmitoylation were obtained with a class imbalance of 6.75:1 in favor of non-*S*-palmitoyl sites (Figure 2A). The dataset was composed of multi- and single-pass TMPs (Figure S1A) with a broad distribution of *S*-palmitoyl sites across cytoplasmic and transmembrane regions (Figure S1B).

**Figure 2.**
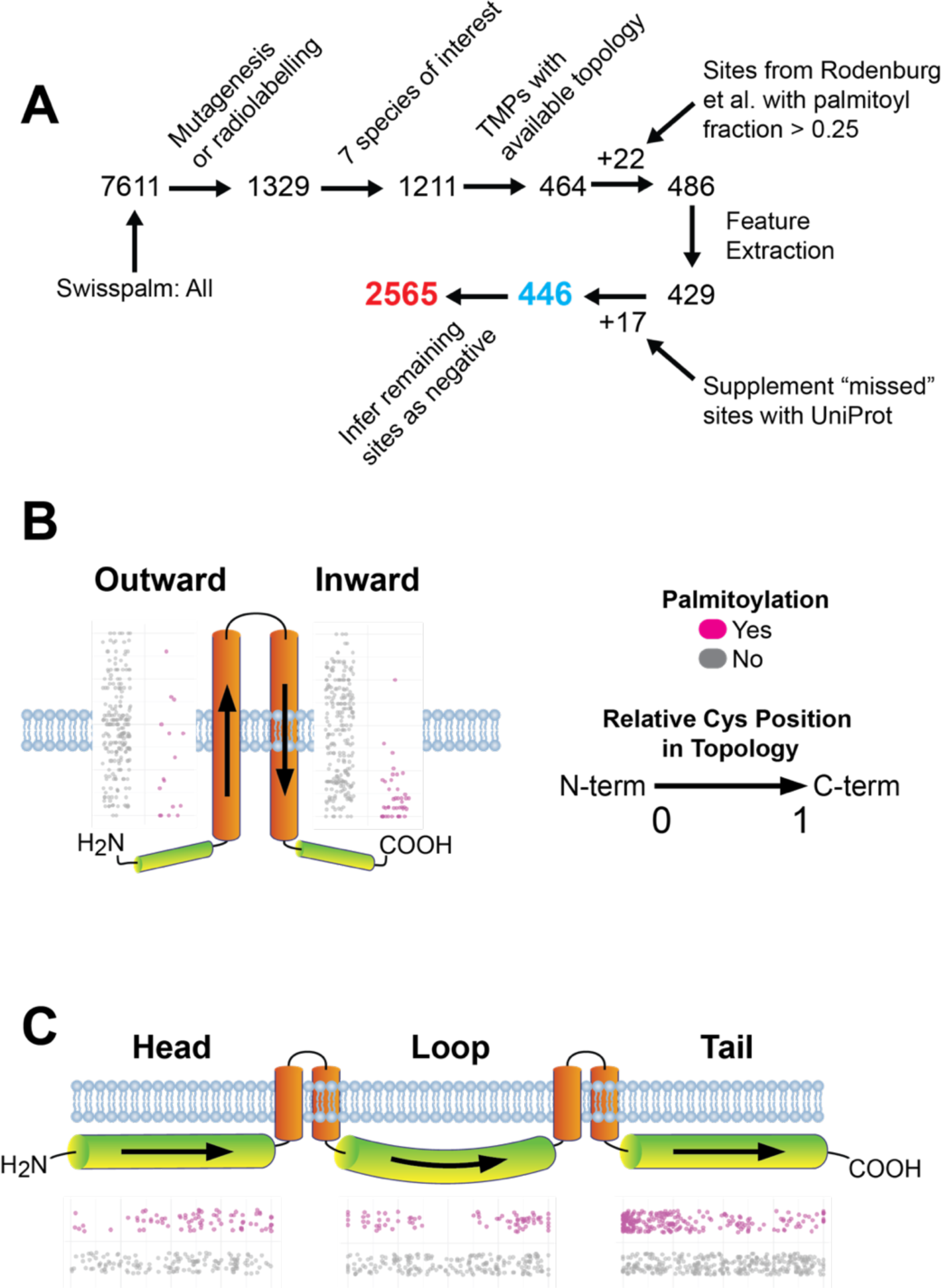
Establishing the training dataset and exploratory data analysis. (A) Schematic of training data isolation and pre-processing. Data were filtered to include sites confirmed by site-directed mutagenesis or ^3^H-palmitate radiolabeling. The seven included species were *H. sapiens* (58.15%), *M. musculus* (23.65%), *R. norvegicus* (11.62%), *A thaliana* (2.59%), *S. cerevisiae* (1.93%), mutant proteins from Rodenburg et al. (0.73%) and *B. taurus* (0.60%). Locations of S-palmitoyl sites within (B) transmembrane and (C) cytoplasmic regions. Shown alongside each region is a jitterplot of relative Cys location in Topology where 0 and 1 represent the N- and C-terminal ends of each region.

### Exploratory data analysis implicates the cytoplasmic-membrane interface in *S*-palmitoylation

Many experiments have confirmed TMP *S*-palmitoylation at juxtamembrane regions(2, 11, 12, 24, 28, 29). To gauge the validity of the training dataset and interrogate the relevance of this proposed feature, the training dataset was analyzed in terms of distance from the cytoplasmic-transmembrane interface. Measuring from the topological end, inward oriented transmembrane domains showed median distances of 1 (mean 2.2) vs. 10 (mean 9.6) amino acids for *S*-palmitoyl vs. non-*S*-palmitoyl sites, respectively (Figure 2B). For outward domains, median distances were 4 (mean 5.8) vs. 10 (mean 10.1) amino acids from the topological start. Consistent with experimental observations, cytoplasmic *S*-palmitoyl sites exhibited a general proximity to the lipid bilayer with median distances of 8.5 and 12 amino acids for cytoplasmic heads and tails, respectively (Figure 2C). In contrast, their non-*S*-palmitoyl Cys counterparts had a median membrane distance of 151 and 116 amino acids for cytoplasmic heads and tails, respectively. Collectively, these measurements from TopoPalmTree’s training data align with the many small-scale observations of *S*-palmitoylation clustering near the transmembrane-cytoplasmic interface.

### *S*-palmitoylation is associated with regional trends in hydrophobicity

Given their juxtamembrane nature, one would expect *S*-palmitoyl hot spots to be in areas with changing hydrophobicity as the polypeptide spans from cytoplasmic to transmembrane regions and vice versa. To capture this biophysical feature, mean hydrophobicity was calculated for each Cys using a window sequence of 5 amino acids in N- and C-terminal directions constituting an absolute (sum) and gradient (difference) in values. A positive gradient represents increasing hydrophobicity towards the C-terminus and vice versa. Within transmembrane domains, total hydrophobicity was higher and lower for *S*-palmitoyl sites on inward and outwards oriented regions, respectively (Figure 2A). Hydrophobicity gradients were more pronounced for *S*-palmitoyl compared to non-*S*-palmitoyl sites, consistent with *S*-palmitoylation clustering at the cytoplasmic-membrane interface where hydrophobicity gradients would be expected to undergo large value swings (Figure S2B). Within cytoplasmic regions, *S*-palmitoyl sites exhibited higher total hydrophobicity (Figure S2C) along with higher gradients (Figure S2).

### Cysteine cooperativity and general feature assessment

Sites of *S*-palmitoylation are often clustered with multiple adjacent Cys residues with evidence of cooperativity(24, 30) or acting as a barrier to de-palmitoylation(31). To build this phenomenon into our feature set, a window Cys scoring scheme was developed that provides points for adjacent Cys residues within the window sequence (Figure S3A). Within the training dataset, *S*-palmitoyl sites showed a higher mean window Cys score (Figure S3B) suggesting that *S*-palmitoyl cooperativity occurs in TMPs and allowing the model to exploit this biological feature.

During exploratory data analysis, a relatively paucity of asparagine residues was noted in the windows of *S*-palmitoyl sites (Figure 3C/D). Interestingly, a similar trend was not observed for glutamine despite these amino acids having structurally similar side chains. Aside from relative asparagine depletion, polybasic residues were readily observed within the windows of cytoplasmic *S*-palmitoyl sites (Figure S4). These are known to promote protein-membrane association(32) and *S*-palmitoylation(33) at least in part through decreasing the Cys thiol pKa(34, 35).

**Figure 3.**
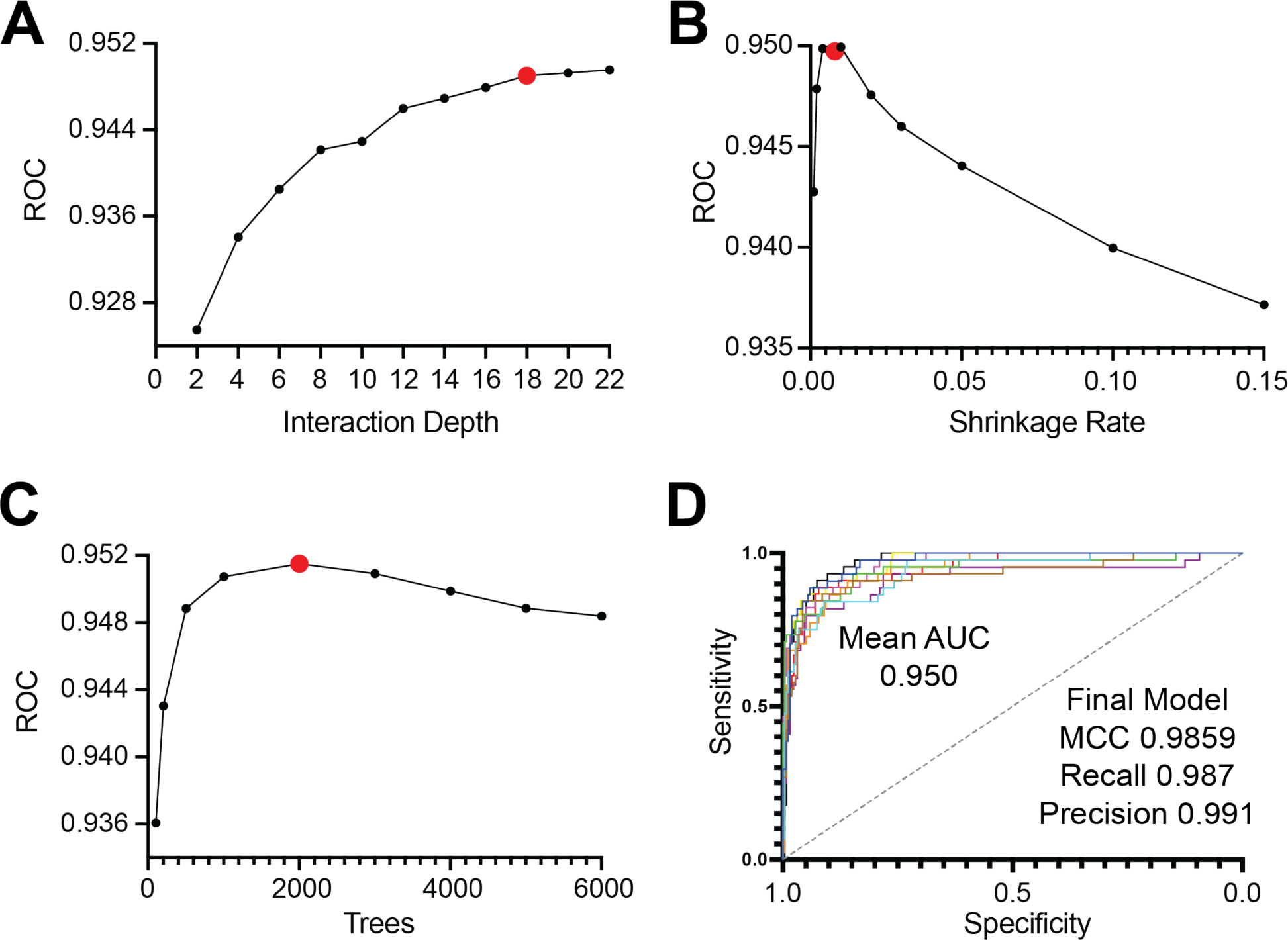
Training and hyperparameter tuning of TopoPalmTree. Shown on each y-axis are mean AUC values from the receiver operator curve vs. (A) interaction depth, (B) number of trees and (C) shrinkage rate. Chosen hyperparameters are highlighted in red. (D) Performance on 10-fold cross validation with chosen hyperparameters of interaction depth = 18, ntree = 2000, shrinkage = 0.08 and m.minobsinode = 5. Statistics relevant to the final model fitting are shown in (D).

### Training the gradient boosted tree and validation with a viral *S*-palmitoyl dataset

The gradient boosted tree (GBT) are a powerful machine learning technique with desirable properties such as model interpretability, flexibility with respect to types of tasks, automatic feature importance, and robust methods to prevent overfitting. Upon creation of 28 distinct features within the training data, TopoPalmTree was trained via the gbm package in R. Hyperparameter tuning was performed through grid search to optimize the interaction depth, shrinkage rate and number of trees (Figure 3A-C). Model performance was relatively robust to hyperparameter choice, exhibiting high AUC on 10-fold cross validation and performance metrics on final model fitting (Figure 3D). Feature importance plotting (Figure S5) confirmed the importance of Cysteine location, Cys score (i.e. cooperativity), hydrophobicity and polarity for the model.

Many viral TMPs rely on *S*-palmitoylation during their infection cycle(4, 36). Given their reliance on host *S*-palmitoyl machinery(37) yet complete lack of sequence homology, viral *S*-palmitoyl TMPs represent an attractive holdout dataset to rigorously assess TopoPalmTree performance. Starting with SwissPalm and experimental criteria of point-mutation or ^3^H-palmitate radiolabeling, 27 viral *S*-palmitoyl TMPs were identified. Another 4 and 5 were identified in UniProt and the primary literature(4), respectively. Of these 36 proteins, 30 had topological annotation available through UniProt. No more than three orthologues were allowed for each type of protein. Following feature extraction, this dataset provided 82 *S*-palmitoyl and 641 non *S*-palmitoyl sites, respectively. Compared to the training dataset, class imbalance of the viral holdout was slightly higher (8.81 vs 6.75).

To avoid the beneficial effects of negative assignments on metrics such as accuracy, the AUC from precision vs. recall was plotted over a continuous threshold range (Figure 4A). The high AUC suggests that either precision or recall can be optimized with relatively little loss in the other metric’s performance. To better understand model performance across all threshold values, the harmonic mean of precision and recall (F1 score) was plotted against threshold (Figure 4B). This plot showed a relatively consistent F1 score of 0.8 suggesting stable performance across the range of thresholds. Utilizing the viral TMP dataset, TopoPalmTree’s performance was compared to GPS-Palm, a recently developed neural network model for *S*-palmitoyl prediction(20) showing even stronger performance than the widely utilized clustering-based tool CSS-Palm(38, 39). Comparing across three threshold values, TopoPalmTree exhibited higher and more consistent Matthews Correlation Coefficient (MCC) than GPS-Palm (Figure 4C). Improving recall via threshold lowering showed that TopoPalmTree’s precision remained relatively preserved, whereas GPS-Palm showed marked deterioration in precision and MCC. Complete results of TopoPalmTree on the viral dataset are listed in Table S2.

**Figure 4.**
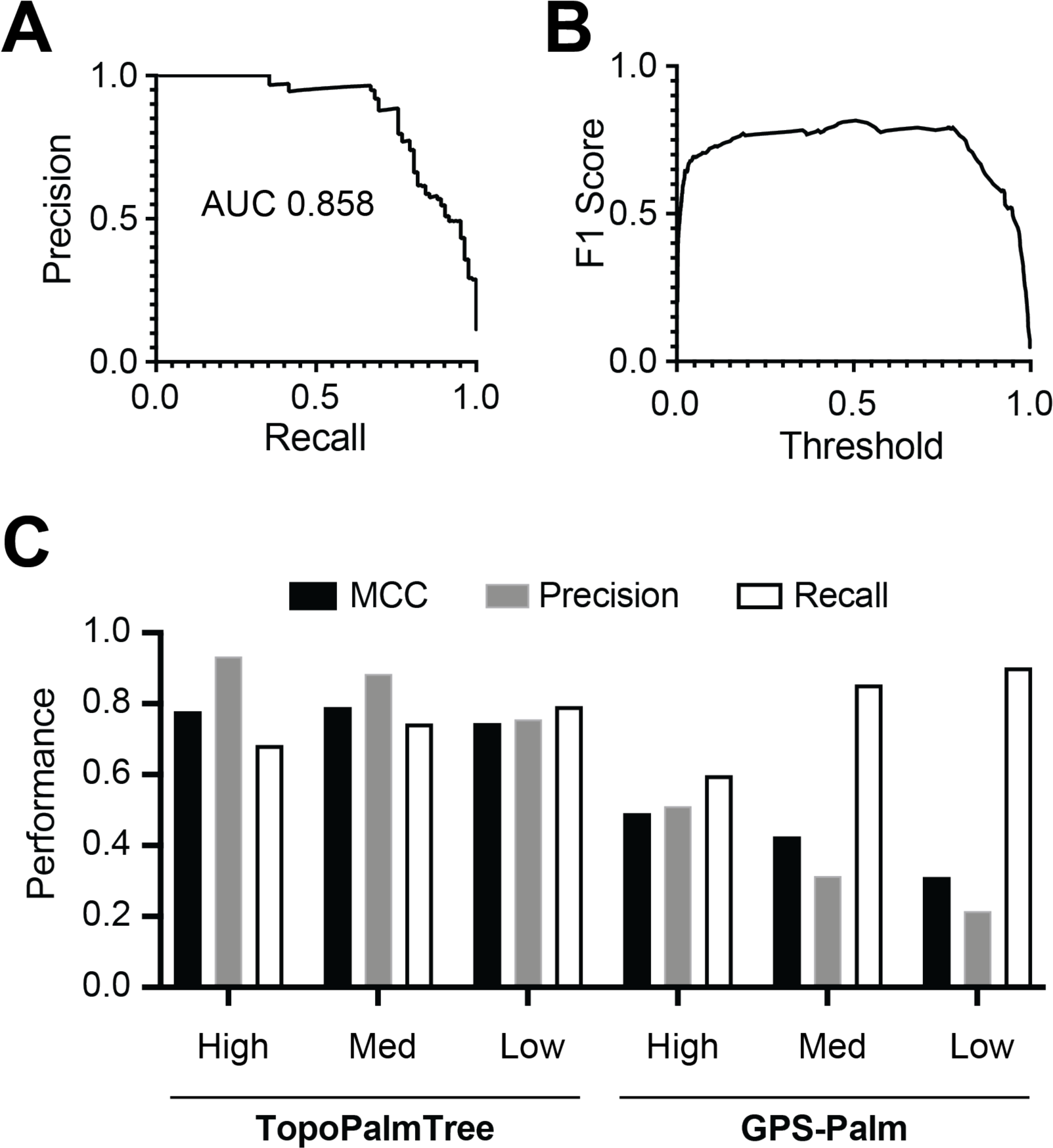
TopoPalmTree validation with a holdout dataset and benchmarking to GPS-Palm. A dataset of 30 viral *S*-palmitoylated proteins (containing 82 *S*-palmitoyl sites) was employed for holdout given complete lack of sequence similarity to training data. Performance shown as (A) precision-recall curve and (B) F1 score vs threshold. (C) Performance of TopoPalmTree at three different thresholds (Low: 0.25, Med: 0.50, High: 0.75) compared to GPS-Palm and the 3 available thresholds of Low, Med, High.

### Application of TopoPalmTree to the murine transmembrane proteome

After filtering the TMP proteome for suitable topological data, 5009 murine TMPs containing 49,828 Cys residues were amenable to inference by TopoPalmTree (Table S1). As shown in Figure 5A, the output from TopoPalmTree revealed 1884 Cys residues (3.8%) deemed high likelihood (probability > 0.75), whereas 45,405 Cys sites (91.1%) have probabilities below 0.25. These findings suggest that TopoPalmTree is discerning towards the positive (*S*-palmitoyl) class and aligns with intuitive expectations of *S*-palmitoylation being restricted to a minority of Cys residues. Of the 1884 putative *S*-palmitoyl Cys sites, 293 are reported (between UniProt and SwissPalm) leaving 1591 sites as novel putative sites of *S*-palmitoylation identified by TopoPalmTree (Figure 5B).

**Figure 5.**
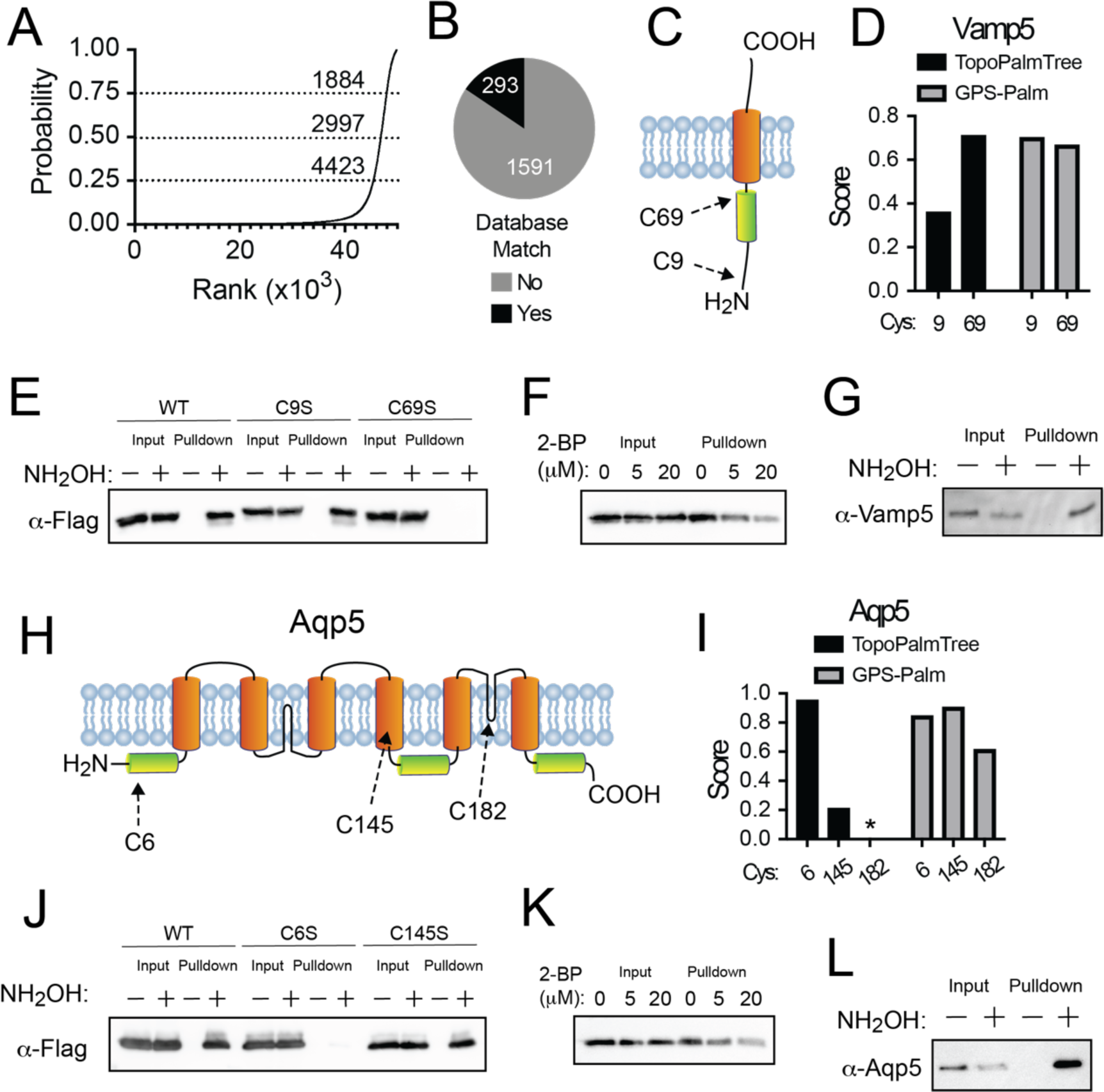
Application of TopoPalmTree for *S*-palmitoyl site discovery. (A) Rank order plot of TopoPalmTree score for each Cys of the murine TMP proteome (49,828 total). Thresholds (dashed lines) are shown at 0.75 (high), 0.50 (medium) and 0.25 (low) with associated number of inferred *S*-palmitoyl sites at each threshold value. (B) Pie chart of database site matches (UniProt and SwissPalm combined) from the high cutoff compared to the number inferred sites that have not been reported in either database. (C) Schematic of Vamp5 topology. (D) Vamp5 probability scores from TopoPalmTree vs GPS-Palm. (E) Acyl-RAC of Vamp5-flag Cys mutants in HEK293 cells. (F) Acyl-RAC of Vamp5-flag from HEK293 cells co-treated with vehicle (DMSO) or 2-BP for 18 h. (G) Acyl-RAC of endogenous Vamp5 in murine lung. (H) Schematic of Aqp5 topology. Shown are transmembrane domains (orange) and two intramembrane regions (black) that do not traverse the entire lipid bilayer. (I) Aqp5 probability scores from TopoPalmTree vs GPS-Palm. Given rare nature of intramembrane regions not represented in the training data, TopoPalmTree does not provide a probability score for Cys^182^ (asterisk). (J) Acyl-RAC of Aqp5-flag Cys mutants in HEK293 cells. (K) Acyl-RAC of Aqp5-flag from HEK293 cells co-treated with DMSO or 2-BP for 18 h. (L) Acyl-RAC of endogenous Aqp5 in murine lung.

### Experimental confirmation and discovery of *S*-palmitoyl sites

To examine the utility of TopoPalmTree to guide discovery of *S*-palmitoylation, two candidate TMPs (Vamp5 and Aquaporin-5) were cloned and subjected to experimental analysis for *S*-acylation. As shown in Figure 5C, Vamp5 has 2 Cys residues: Cys9 located near the N-terminus and juxtamembrane Cys69 located 4 amino acids from the transmembrane domain. As shown in Figure 5D, TopoPalmTree showed higher scoring for Cys69 compared to Cys9 (0.71 vs 0.36), while GPS-Palm exhibited slightly higher score for Cys9 than Cys69 (0.70 vs 0.67). When subjected to experimental analysis for *S*-acylation by Acyl-RAC(40), Vamp5-transfected HEK293 cells exhibited hydroxylamine-dependent pulldown that was completely lost upon mutation of Cys69 to Serine (Figure 5E). Pulldown was dose-dependently inhibited 2-bromopalmitate (Figure 5F) and observed endogenously in murine lung (Figure FG). Notably, Cys69 is flanked by Arg residues and is essentially “lost” as a dipeptide when subjected to trypsinization. This likely explains why Vamp5 Cys69 has never been reported as a site of *S*-palmitoylation, yet it was readily identified by TopoPalmTree. Collectively these findings establish Cys69 as the site of *S*-palmitoylation on Vamp5 and demonstrate the utility of TopoPalmTree in identifying *S*-palmitoylation sites, particularly those that may be missed by bottom-up proteomics.

Another candidate *S*-palmitoyl TMP is Aquaporin-5 (Aqp5), a multi-pass TMP enriched in alveolar type 1 cells and is responsible for water transport across the alveolar membrane(41, 42). Aquaporin 5 has three Cys residues: cytoplasmic Cys6 located 7 amino acids from the lipid bilayer, Cys145 located within a transmembrane region and Cys182 located in a so-called “intramembrane” region buried inside the lipid bilayer yet not spanning the membrane (Figure 5H). These intramembrane domains are present within only 1.1% of the murine proteome. Given their paucity and lack of representation in the training dataset, intramembrane Cys residues (e.g. Aqp5 Cys182) were omitted from inference by TopoPalmTree. As shown in Figure 6I, TopoPalmTree strongly favored Cys6 over Cys145 (score 0.95 vs 0.21). In contrast, GPS-Palm suggested slight preference for Cys145 (0.90) followed by Cys6 (0.84) and Cys182 (0.61). Of note, Aqp5 Cys6 was detected in a proteomic study of bovine lens(43) but has not been subjected to prediction methods or confirmatory studies such as site-directed mutagenesis. As shown in Figure 6J, HEK293 cells transfected with Aqp5 showed hydroxylamine-dependent pulldown by the Acyl-RAC assay with loss of signal upon mutation of Cys6 to Ser. Mutation of Cys145 to Ser had no effect, again demonstrating superior performance of TopoPalmTree in identifying the correct site of *S*-palmitoylation. Pulldown of Aqp5 was dose dependently inhibited by 2-bromopalmitate (Figure 6K) and observed endogenously in murine lung (Figure 5L). Similar to Vamp5, Aqp5 serves as another proof-of-concept that TopoPalmTree can recognize sites of TMP *S*-palmitoylation and facilitate targeted confirmatory experiments.

**Figure 6.**
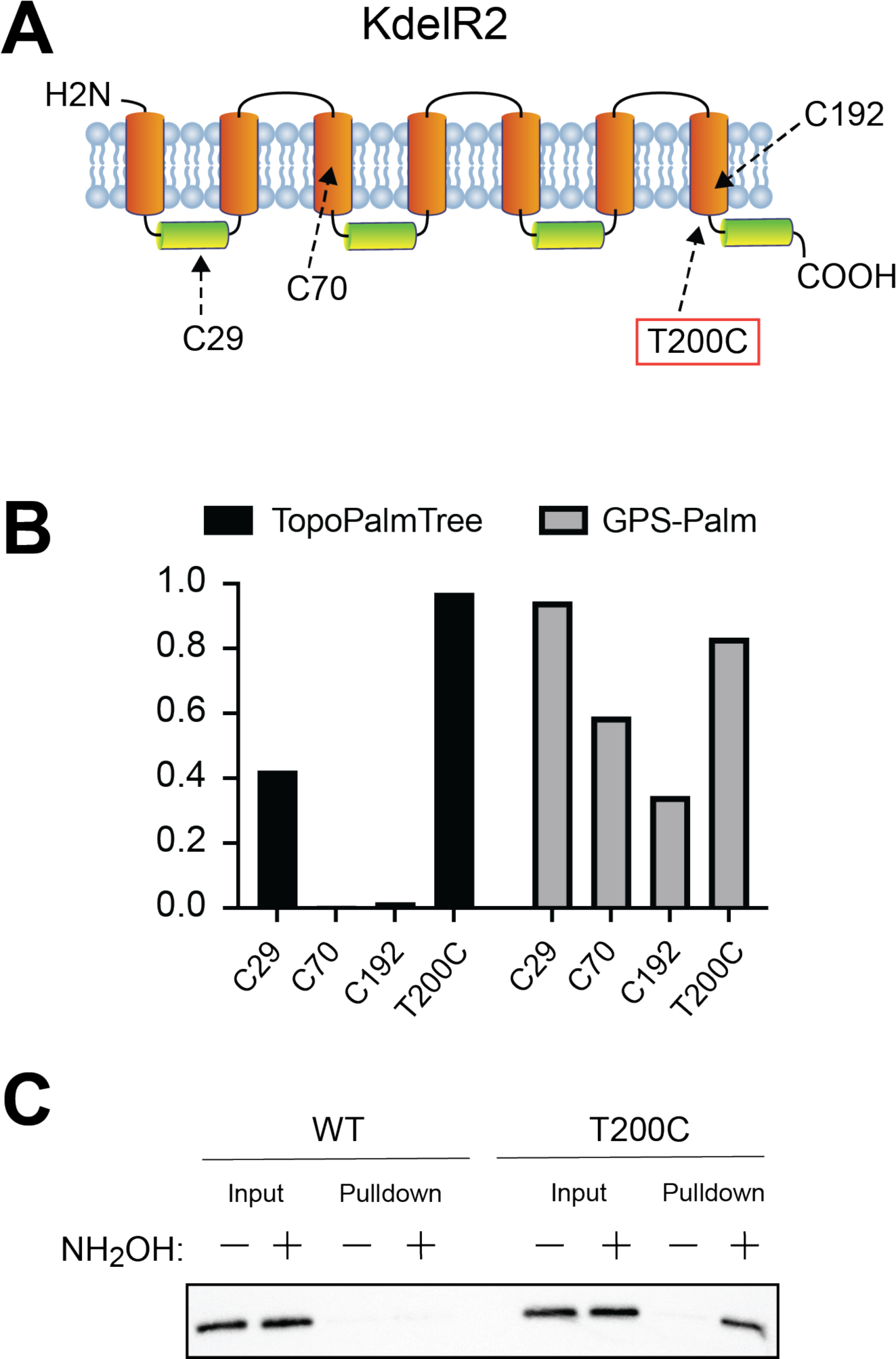
TopoPalmTree facilitates in silico “design” of an *S*-palmitoyl site. (A) Schematic of KdelR2 along with (B) probability scores of its 3 native Cys residues. Shown in red is position 200, which is located in the C-terminal juxtamembrane region where 9 residues showed probability scores > 0.90 by *in silico* mutagenesis. (C) Acyl-RAC of WT vs T200C KdelR2-flag in HEK293 cells. Hydroxylamine-dependent pulldown is indicative of protein *S*-acylation status.

### Rational design of an *S*-palmitoyl site on KdelR2

Having established the utility of TopoPalmTree to identify sites of *S*-palmitoylation, we next questioned whether TopoPalmTree could be used to design a site of *S*-palmitoylation into a protein that is otherwise not *S*-palmitoylated. To identify a protein with Cys residues that was presumably not *S*-palmitoylated, we first screened for proteins that had at least 3 Cys residues with probability scores < 0.5 for each. One candidate KDEL Receptor 2 (KdelR2) – a TMP that facilitates retrieval proteins from the Golgi back to the ER – has 3 Cys residues (Figure 6A) with TopoPalmTree generating very low probabilities for Cys70 and Cys192 (Figure 6B) both of which are located within transmembrane domains. TopoPalmTree provides a low-moderate score for Cys29 (0.42) located in the middle of a cytoplasmic loop. On the contrary, GPS-Palm indicates a high score for Cys29 (0.94) with decreasing scores for Cys70 (0.59) and Cys192 (0.35). When subjected to *in silico* Cys-scanning mutagenesis followed by TopoPalmTree inference, 9 of the 11 locations with probability scores > 0.90 were located on the C-terminal juxtamembrane region (residues 200 – 205) and adjacent C-terminus (residues 210-212). One candidate mutant (KdelR2 T200C) was subjected to Acyl-RAC alongside wild-type KdelR2. As shown in Figure 6C, the incorporation of a Cys residue at position 200 results in hydroxylamine-dependent pulldown of KdelR2, thus confirming the ability of TopoPalmTree to guide the rational design of an *S*-palmitoyl site.

## DISCUSSION

Protein *S*-palmitoylation is a prevalent hydrophobic modification that imparts hydrophobicity and regulates a diverse array of biological processes. Small-scale studies have consistently observed *S*-palmitoylation at juxtamembrane residues of TMPs, and our findings suggest these sites are incompletely characterized by current experimental techniques due to their inherently hydrophobic locations. To address this methodological gap, a GBT based on simple UniProt-derived features (TopoPalmTree) was created from curated *S*-palmitoyl data, validated on an unrelated dataset of viral *S*-palmitoyl proteins to rigorously assess feature meaningfulness, experimentally applied to identify sites of *S*-palmitoylation and subject a TMP (KdelR2) to *S*-palmitoyl rational design.

Unlike exceedingly complex algorithms, TopoPalmTree uses a relatively straightforward model – gradient boosted trees – requiring very little computing resources and can be trained in an R environment such as RStudio. Being derived from UniProt, the feature set is readily accessible. Thus, the described topological concepts can be easily implemented and adapted for future *in silico S*-palmitoylation studies and facilitate further expansion of the *S*-palmitoyl toolbox. For example, future directions for TopoPalmTree could include merging with chemoproteomic or alternative PTM datasets to understand the interplay of *S*-palmitoylation with drug binding or other PTMs, respectively.

While the utility of TopoPalmTree has been demonstrated, the approach has several significant limitations. Firstly, TopoPalmTree is restricted to TMPs as the utilized topological feature set is not applicable to soluble (or membrane-associated) proteins. Secondly, TopoPalmTree’s reliance on UniProt-derived topological information means that the model is limited to the number of proteins with available topological data. In the case of the murine proteome, approximately one quarter of TMPs (1065 total) have incomplete annotation leaving a significant portion of the TMP proteome inaccessible to TopoPalmTree. This limitation can be addressed with continued topological annotation of TMPs. Despite these limitations, TopoPalmTree should provide a valuable resource for future studies on TMP *S*-palmitoylation, particularly those focused on the molecular and functional consequences of this increasingly appreciated post-translational modification.

## EXPERIMENTAL PROCEDURES

### Reagents and Cell Culture

All chemicals and reagents specific to cloning and cell culture are listed in table format in the supporting information. HEK293 cells were cultured in DMEM supplemented with 10% FBS and 100 U/ml penicillin-streptomycin. Cells were grown in a 5% CO2 atmosphere. Transfections were performed for 18-24 h using 2.5:1 (µl: µg) ratio polyethyleneimine (PEI) to plasmid in OptiMem (50 µl per 1 µg plasmid). For most experiments, a 6 cm TC-treated plate was transfected with 3 µg of indicated plasmid. SDS-PAGE was performed in 12% acrylamide using the Bio-Rad Mini-Protean System. Transfer to PVDF was performed with the Trans-Blot SD Semi Dry Transfer Cell (Bio-Rad). Membrane blocking, primary and secondary antibody exposures were performed in TBST containing 5% dried milk (w/v). Primary antibody was incubated overnight. Membranes were visualized with Clarity Western ECL Substrate (Bio-Rad) and a Chemidoc MP imager (Bio-Rad).

### Biophysical characterization of juxtamembrane and swisspalm derived *S*-palmitoyl sites

The murine proteome from UniProtKB was read into RStudio (version 2023.12.1+402). Each cysteine was extracted from the protein sequence into a new row followed by in silico trypsinization on the C-terminal side of every Lys or Arg residue except when followed by Pro. All Cys-containing peptides subjected to molecular weight and mean hydrophobicity measurements based on Kyte-Doolittle scale(44). Visualizations were performed using ggplot2 and Prism.

### Training dataset and gradient boosted tree

The complete dataset of “Sites” was downloaded from Swisspalm (Version 2022-09-03) and filtered within column “site_techniques” for terms “Point mutation” or “palmitate”. Upon filtering for the 7 indicated species and addition of 22 positive class data corresponding to all sites with palmitoyl fraction > 0.25 from Rodenburg et al.(24), stringr package was employed to split topological data into new columns along with character to numeric class conversions. Topological lengths along with relative and absolute distances from the ends of each topology were calculated for every Cys residue. Less common topological locations such as “Lumenal” and “Stromal” were reclassified as “Extracellular” to ensure adequate alignment of categorical variables. The orientation of transmembrane domains (inward vs outward) was determined by identifying a row that 1) shared the same accession number and 2) had a stop position that was 1 integer lower than the transmembrane’s start position. If the preceding domain was identified as “extracellular/lumenal” or “cytoplasmic” the transmembrane domain was considered inward or outward, respectively. Five amino acid windows were extracted on the N- and C-terminal side of each Cys residue along with calculation of Cys scoring, hydrophobicity, charge states, polarity, aliphatic index, and transmembrane tendency were calculated and each assigned its own column. To ensure complete assignment of the positive class, the SwissPalm-derived *S*-palmitoyl assignments were cross-referenced to UniProt leading to the re-assignment of 17 sites from negative to positive. Upon establishment of the training dataset, the gbm package was utilized for implementation of a gradient boosted machine algorithm. Training was configured via the caret package. Control parameters included: method = “cv”, number = 10, classProbs = TRUE and summaryFunction = twoClassSummary. After tuning by interaction grid, final hyperparameters were interaction.depth = 18, n.trees = 2000, shrinkage = 0.008 and n.minobsinnode = 5.

### Viral validation dataset

The viral holdout dataset was assembled from 3 sources with no more than 3 orthologues per protein type: 27 proteins from UniProtKB, 5 proteins from the primary literature summarized by Veit(4), and 4 proteins from SwissPalm verified by radiolabeling or site-directed mutagenesis. Of these 36 proteins, 30 had suitable topographical annotations from UniProt. Following feature extraction, the model was applied to the dataset with probability thresholds of 0.25, 0.5 and 0.75 for binary classification. The same data were exported in FASTA format, submitted to Windows format GPS-Palm in batch format and read back into Rstudio for analysis.

### Cloning and site directed mutagenesis

To obtain a cDNA library, RNA was isolated from murine lung with the RNAeasy kit (Qiagen) followed by reverse transcription with the SuperScript III Reverse Transcription kit (Invitrogen) using oligo-dT primers. DNA’s were amplified by PCR and subjected to restriction-ligation into pCMV-EGFP. All clones were confirmed by Sanger sequencing. Site directed mutagenesis was performed using the NEBasechanger tool and NEB’s Q5 mutagenesis protocol. All relevant oligonucleotide sequences are listed in the supporting information.

### Annealed oligo ligation

To replace the EGFP with a flag epitope tag, two micrograms of EGFP-containing plasmid was subjected to digestion with AgeI and Not I and gel extracted using the Qiagen Qiaex II kit. The flag-encoding oligos (100 µM each) were subjected to phosphorylation with T4 PNK according to the NEB protocol at 37C for 30 min. After heating to 95 C for 5 min, annealing was accomplished with ramp to 25 C at 5 C/min. The phosphorylated and annealed oligo was diluted 1:100 into nuclease free water, then 1 µL added to a 10 µL T4 ligation reaction including 50 ng of digested vector. After 30 min at room temp, 1 µL of ligation was used to transform chemically competent E coli. Colonies were confirmed by Sanger sequencing.

### Assay of *S*-palmitoylation by Acyl-RAC

The Acyl-RAC assay was performed as described(40) with several modifications. For each sample, one 6 cm dish of HEK293 cells or 1 mg of murine lung tissue was subjected to lysis in 100 mM HEPES, 5 mM EDTA, 0.5% Triton X-100, pH 7.2 containing 20 mM N-ethyl maleimide (NEM, made from a fresh 1M stock solution in MeOH). Following probe sonication on ice, lysates were centrifuged at 5000g for 10 min and supernatant transferred into 2 ml blocking reactions containing 100 mM HEPES, 5 mM EDTA, 1% SDS and 20 mM NEM. Blocking was performed at 50 C for 1 hour with frequent vortexing, then proteins were precipitated with 3 volumes (6 ml of room temperature MeOH). Samples were mixed, incubated at −20 C for 30 min and pellets recovered with centrifugation at 3000g for 5 min. Liquid was aspirated, and the dried pellet was resuspended in 10 ml of MeOH by vortexing (this step ensures complete removal of NEM from the blocking step, which can compete with capture). Protein was again collected with centrifugation at 3000g for 5 min, dried and resuspended in 800 µl of 100 mM HEPES, 5 mM EDTA, 1% SDS pH 7.2. Half of each reaction was transferred onto 25 µl of pyridyl disulfide sepharose (PDS) followed by addition of either water or 2M neutral NH_2_OH to final concentration of 500 mM NH_2_OH. Samples were rotated at room temperature for 12-18 h, washed three times with 1 ml of wash buffer (50 mM HEPES, 2 mM EDTA, 1 % SDS) and eluted with 80 µl of wash buffer containing 20 mM DTT. After 20 minutes at room temperature, eluant was collected, mixed with 4x Laemlli buffer, heated to 95C for 5 min then analyzed by SDS-PAGE with western blotting.

## DATA AVAILABILITY

All original code is available at https://drive.google.com/drive/folders/1FAfElxsRTymUac68VN2UaIT0BWoTy_X1?usp=drive_link Please see README.txt for information regarding R markdown and associated csv files

## SUPPORTING INFORMATION

This article contains supporting information.

## ACKNOWLEDGEMENTS

We thank Patrick J. Casey (Duke Pharmacology and Cancer Biology), Christine E. Eyler (Duke Radiation Oncology) and Cliburn Chan (Duke Biostatistics & Bioinformatics) for helpful advice.

## AUTHOR CONTRIBUTIONS (CRediT STATEMENT)

MTF: Conceptualization, Methodology, Software, Validation, Investigation, Data Curation, Writing - Original Draft, Visualization, Supervision, Funding acquisition; JRE.: Investigation, Writing - Review & Editing; SO: Conceptualization, Formal analysis, Visualization, Writing - Review & Editing; RS: Formal analysis, Supervision, Writing - Review & Editing; PRT: Conceptualization, Resources, Supervision, Writing - Review & Editing, Funding acquisition.

## FUNDING AND ADDITIONAL INFORMATION

The work was supported in part by an award from the Duke Office of Physician Scientist Development (OPSD) and T32HL160494 (to M.T.F.) and research awards from NHLBI/NIH (R01HL160939, and R01HL153375) to P.R.T.

