## Supporting Information for "Topology-Driven Discovery of Transmembrane Protein *S*-Palmitoylation"

TABLE OF CONTENTS

Supplementary methods 2

Figure S1. Summary statistics of the training dataset 5

Figure S2. Hydrophobicity related features 6

Figure S3. Amino-acid specific features 7

Figure S4. Sequence logos based on *S*-palmitoyl status 8

Figure S5. Feature importance plot for TopoPalmTree. 9

Supplementary Table Descriptions 10

Table S1. TopoPalmTree applied to viral *S*-palmitoyl protein dataset (xls)

Table S2. TopoPalmTree applied to the mouse transmembrane proteome (xls)

SUPPLEMENTARY METHODS

*Synthesis of pyridyl disulfide sepharose for Acyl-RAC* – All reactions are conducted in PDS buffer (100 mM HEPES + 2 mM EDTA pH 7.8) unless otherwise indicated. A 3 mL slurry of NHS-Sepharose was washed three times with PDS buffer in a 10 mL filter column, followed by addition of 100 mM cystamine in PDS buffer to fill the 10 ml column. The tube is rotated for 4 hours at room temperature, washed at least 4 times with PDS buffer, washed 4 times with PDS buffer containing 20 mM DTT, then washed another 4 times with PDS without DTT. The slurry is washed 4 times with MeOH, 100 mM 2-pyridyl disulfide (aka Aldrithiol-2) in MeOH is added to fill the column and the reaction is rotated at room temperature for 15 minutes. The slurry is washed twice with 100 mM 2-pyridyl disulfide in MeOH then three times with MeOH, then finally isopropanol. The slurry is stored at 50 % density (e.g. 2 ml of sepharose in 4 ml of isopropanol) in a sterile 15 ml conical tube at 4°C.

**Resource Table**

| REAGENT or RESOURCE | SOURCE | IDENTIFIER |
|---|---|---|
| **Antibodies (dilution)** | | |
| Flag M2 monoclonal Ab (1:4000) | Sigma | F3165 |
| Flag monoclonal Ab (1:4000) | Proteintech | 66008-4-Ig |
| Vamp5 polyclonal Ab (1:1000) | Proteintech | 11822-1-AP |
| Aqp5 polyclonal Ab (1:1000) | Cell Signaling Technologies | 59558 |
| **Reagents for synthesis of thiopropyl sepharose (TPS)** | | |
| Sepharose-NHS | Sigma | H8280 |
| Cystamine-HCl | Sigma | C121509 |
| Alrithiol-2 (2-PDS) | Sigma | 143049 |
| Pierce Centrifuge Columns, 10 mL | Pierce | 89898 |
| **Enzymes** | | |
| Q5 Polymerase | New England Biolabs | M0491 |
| T4 Ligase | New England Biolabs | M0202 |
| Kinase / Ligase / Dpn1 (KLD Mix) | New England Biolabs | M0554S |
| AgeI-HF | New England Biolabs | R3552 |
| EcoRI-HF | New England Biolabs | R3101 |
| NotI-HF | New England Biolabs | R3189 |

| | | |
|---|---|---|
| **Kits for cloning** | | |
| RNeasy | Qiagen | 74104 |
| Qiaex II Gel Extraction Kit | Qiagen | 20021 |
| SuperScript III First Strand Kit | Thermo | 18080-400 |
| **Cell Lines and Culture Reagents** | | |
| HEK293 | Duke Cell Culture Facility | N/A |
| High Glucose DMEM | Gibco / Thermo Fisher | 11965092 |
| FBS | Cytiva Hyclone | SH30071 |
| Pen-Strep | | |
| Polyethylenimine (PEI) | Polysciences | 23966 |
| Trypsin-EDTA (0.25%) | Gibco / Thermo Fisher | 25200056 |
| **Mammalian expression vector** | | |
| pCMV-EGFP | Addgene | 11153 |
| **Oligonucleotides for cloning and annealed oligo ligation** | | |
| See DNA oligos table below | | |
| **Chemicals / General Reagents** | | |
| 2-Bromohexadecanoic acid (2-bromopalmitate) | Sigma | 238422 |
| N-Hydroxysuccinimidyl-Sepharose™ 4 Fast Flow | Sigma | H8280 |
| 2-Pyridyl disulfide (Aldrithiol-2) | Sigma | 143049 |
| Cystamine dihydrochloride | Sigma | C121509 |

*DNA oligos used for PCR and cloning* – Sequences are all for murine genes. Fw = forward. Rv = reverse. Tm = melting temperature. Ta = annealing temperature used for PCR. For all reactions, Q5 polymerase was used except for Vamp5 cloning into pCMV-EGFP which was performed with Dream Taq.

| Primer Name | Sequence (5’ -> 3’) | Tm / Ta (C) |
|---|---|---|
| **Cloning into pCMV-EGFP** | | |
| Vamp5_EcoRI_Fwd | ATTAGAATTCACCATGGCAGGGAAAGAACTGAAGCAATGCCAGC | 70 / 66 |

| | | |
|---|---|---|
| Vamp5_AgeI_Rev | ATTAACCGGTGGTTTACTACTGTCCCCACCACTCGGAAG | 71 / 66 |
| Aqp5_EcoRI_Fwd | TATAGAATTCACCATGAAGAAGGAGGTGTGTTCAGTTGC | 65 / 68 |
| Aqp5_AgeI_Rev | TAATACCGGTAAGTGTGCCGTCAGCTCG | 67 / 68 |
| KdelR2_EcoRI_Fwd | ATTAGAATTCACCATGAACATCTTCCGGCTGACTGGGGACCTG | 70 / 72 |
| Kdelr2_AgeI_Rev | TTAAACCGGTGCTGGCAGGCTGAGCTTCTTCCCTTTGAGTAC | 73 / 72 |
| **Site Directed Mutagenesis** | | |
| Vamp5_C9S_Fwd | ACTGAAGCAAAGCCAGCAGCAGG | 70 / 66 |
| Vamp5_C9S_Rev | TCTTTCCCTGCCATGGTG | 65 / 66 |
| Vamp5_C69S_Fwd | AAATATCCGGAGCCGGGTCTACT | 67 / 67 |
| Vamp5_C69S_Rev | TCCCAGCGCTTCTGC | 66 / 67 |
| Aqp5_C6S_Fwd | GAAGGAGGTGAGCTCAGTTGCCT | 64 / 61 |
| Aqp5_C6S_Rev | TTCATGGTGAATTCGAAGC | 60 / 61 |
| Aqp5_C145S_Fwd | GCTGGCCCTCAGCATCTTCTCCT | 70 / 66 |
| Aqp5_C145S_Rev | TTCATGGTGAATTCGAAGC | 65 / 66 |
| KdelR2_T200C_Fwd | CTTGTACATTTGCAAAGTACTCAAAGGGAAGAAG | 57 / 58 |
| KdelR2_T200C_Rev | TAGAAGAAGTCGCAGTAG | 58 / 58 |
| **Annealed oligos for EGFP removal and flag epitope insertion at Age I / Not I sites** | | |
| EGFP_to_flag_Fwd | CCGGTAGACTACAAAGACGATGACGACAAGTGATCTAGAGC | N/A |
| EGFP_to_flag_Rev | GGCCGCTCTAGATCACTTGTCGTCATCGTCTTTGTAGTCTA | N/A |

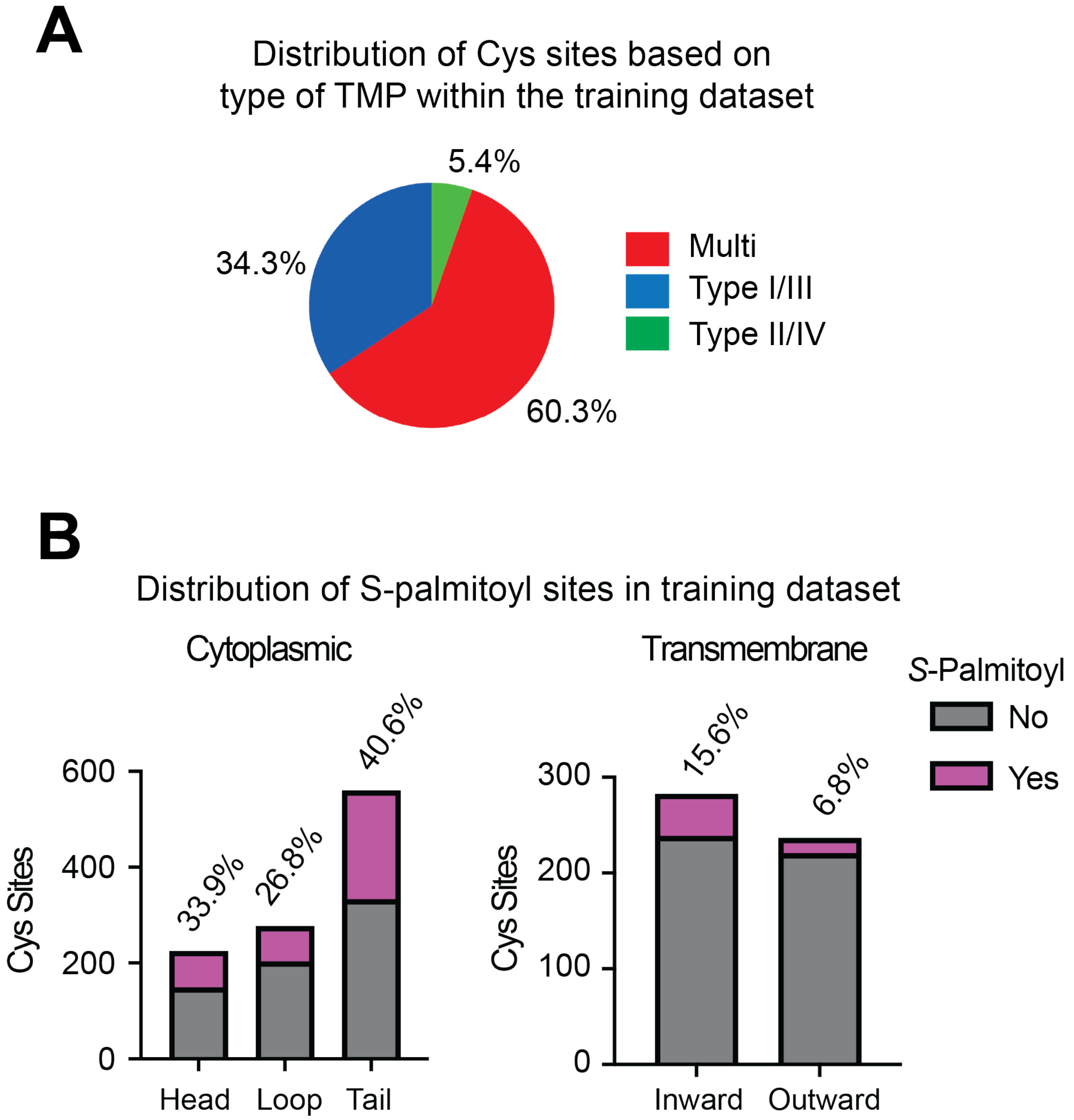

**Figure S1. Summary statistics of the training dataset.** (A) Types of TMPs represented in the training data. Total number of Cys sites are grouped based on whether they reside on a multi-pass, Type I/III or Type II/IV TMP (based on orientation of the N-termini relative to the lipid bilayer). (B) Distribution of *S*-palmitoyl (magenta) and non-*S*-palmitoyl (gray) Cys sites based on topological location. Shown above each column are the percentages of Cys sites that are *S*-palmitoylated within each type.

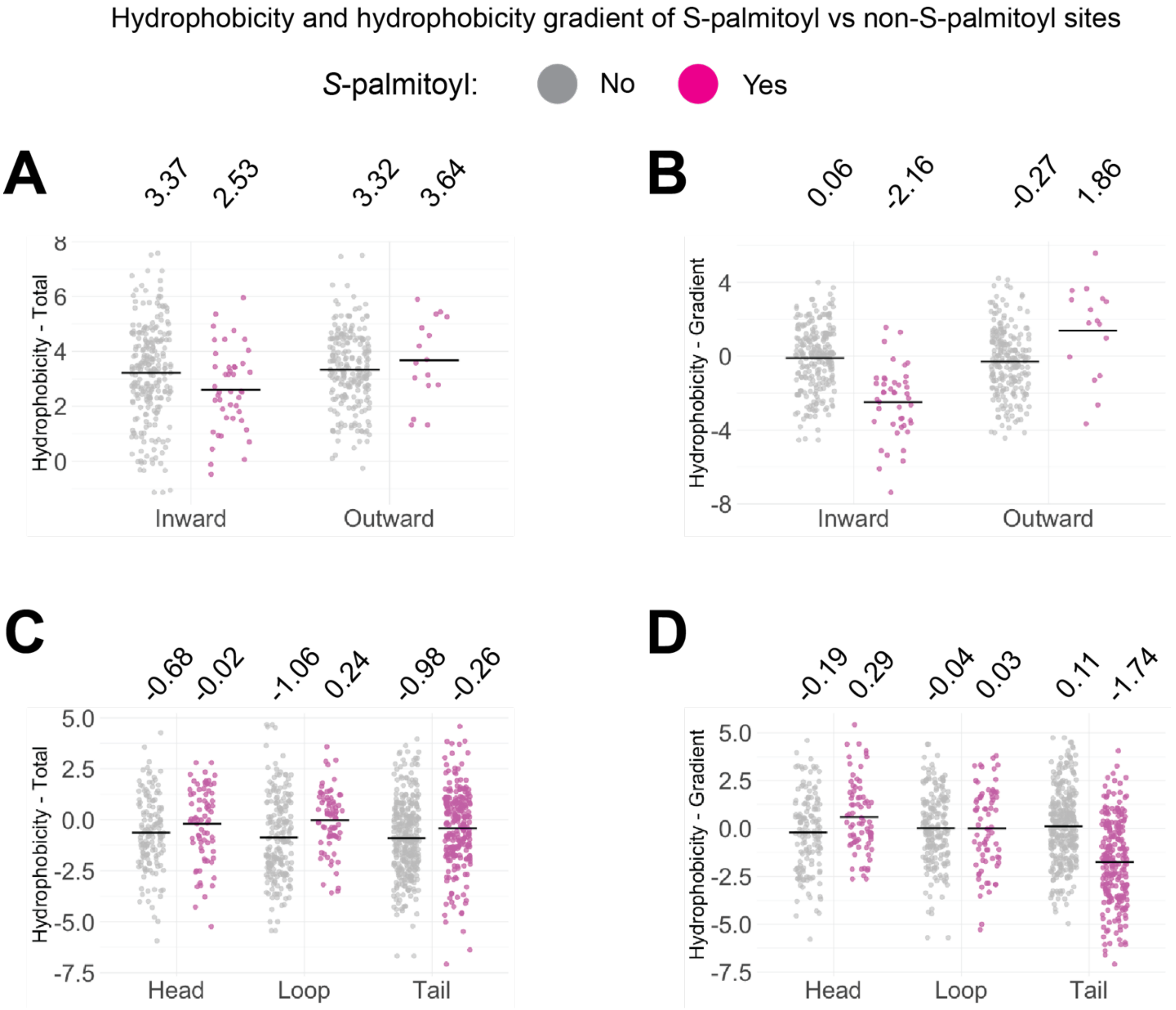

**Figure S2. Hydrophobicity and hydrophobicity gradients as examples of the TopoPalmTree feature set.** Hydrophobicity values are derived from the window sequence (5 amino acids flanking on each side of the given Cys residue). Total hydrophobicity is calculated for the entire window sequence whereas the hydrophobicity gradient is the difference in hydrophobicity (C-terminal minus N-terminal window). For total hydrophobicity, positive values indicate increased overall hydrophobicity surrounding the Cys residue. For hydrophobicity gradient, more positive values indicate increasing hydrophobicity from the N- to C-terminal direction across the given Cys residue. Shown are (A) total hydrophobicity and (B) hydrophobicity gradient for transmembrane regions, as well as (C) total hydrophobicity and (D) hydrophobicity gradient for cytoplasmic regions. The black bars represent mean values.

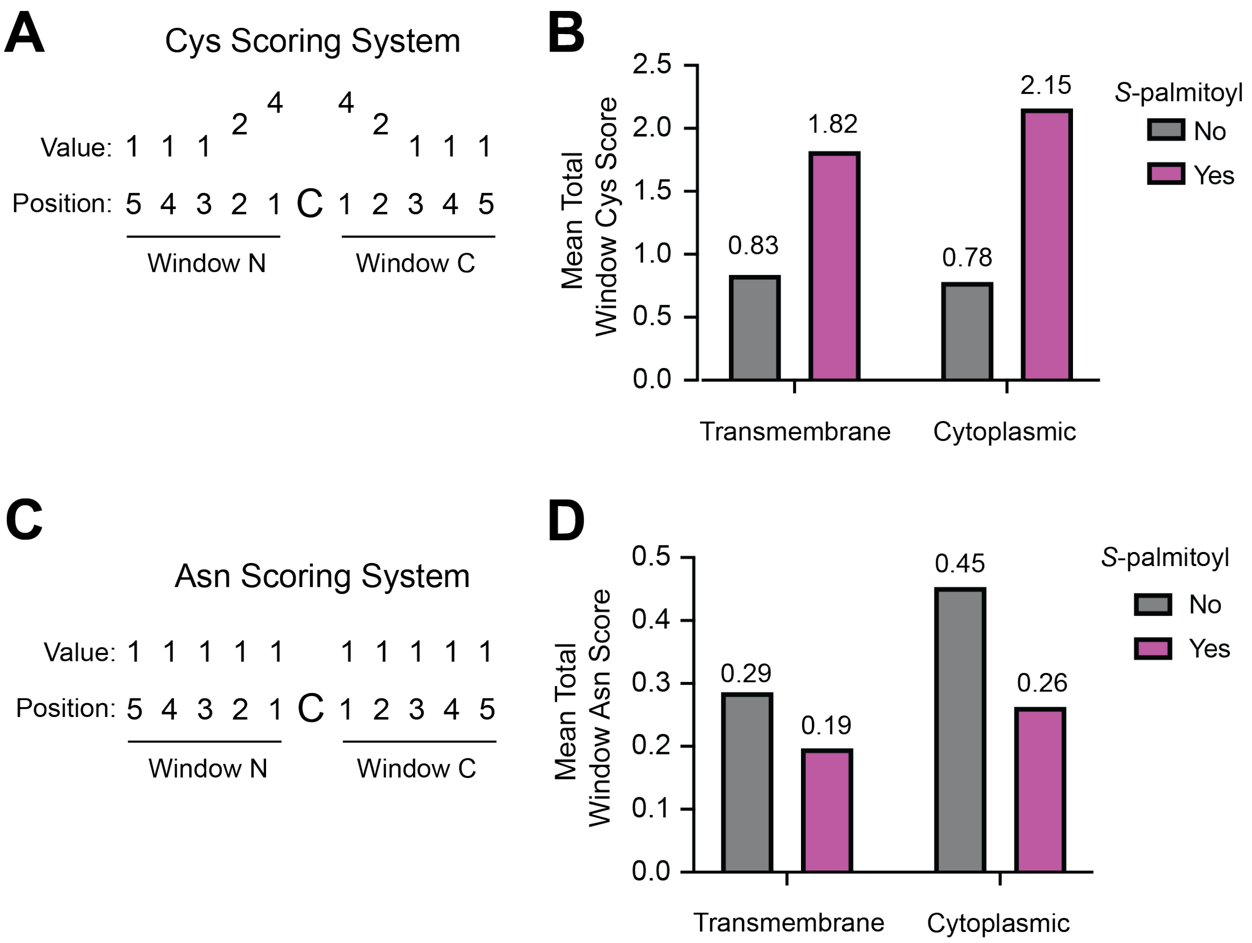

**Figure S3. Window Cys score and Asn scores as examples of amino acid-based features within the training dataset.** (A) Window sequence scoring system for adjacent Cys residues giving priority to closer Cys residue. Cys residues flanking the Cys of interest (center) are awarded 4 points. Cys residues in the 2nd flanking positions are given 2 points, and then 1 point for any Cys within flanking positions 3 through 5 in either direction. (B) The values are then summed to achieve a total score for the N- and C-terminal windows, with mean values shown based on *S*-palmitoyl status. (C) Asparagine scoring system providing one point for every Asn residue within the N- and C-terminal windows. (D) Mean total Asn scores based on *S*-palmitoyl status.

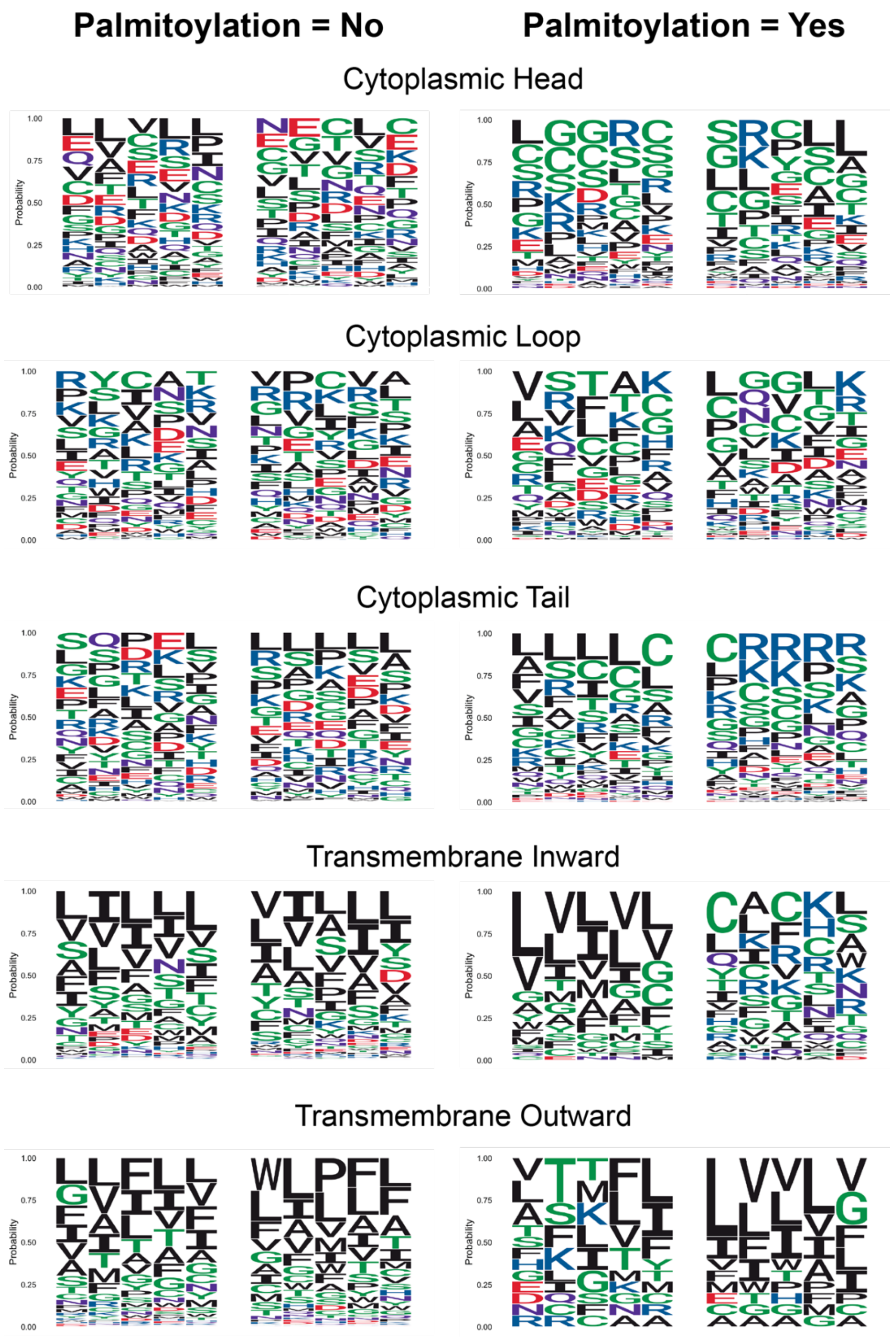

**Figure S4. Sequence logos based on *S*-palmitoyl status within the training dataset.** Sequence logos were generated with package ggseqlogo. For each logo, the central empty space represents the Cys site of interest with N- and C-terminal flanking residue probabilities on the left and right, respectively. The y-axes represent probability with each position summing to 1. Logos are separated by column for *S*-palmitoyl status and row for topological locations.

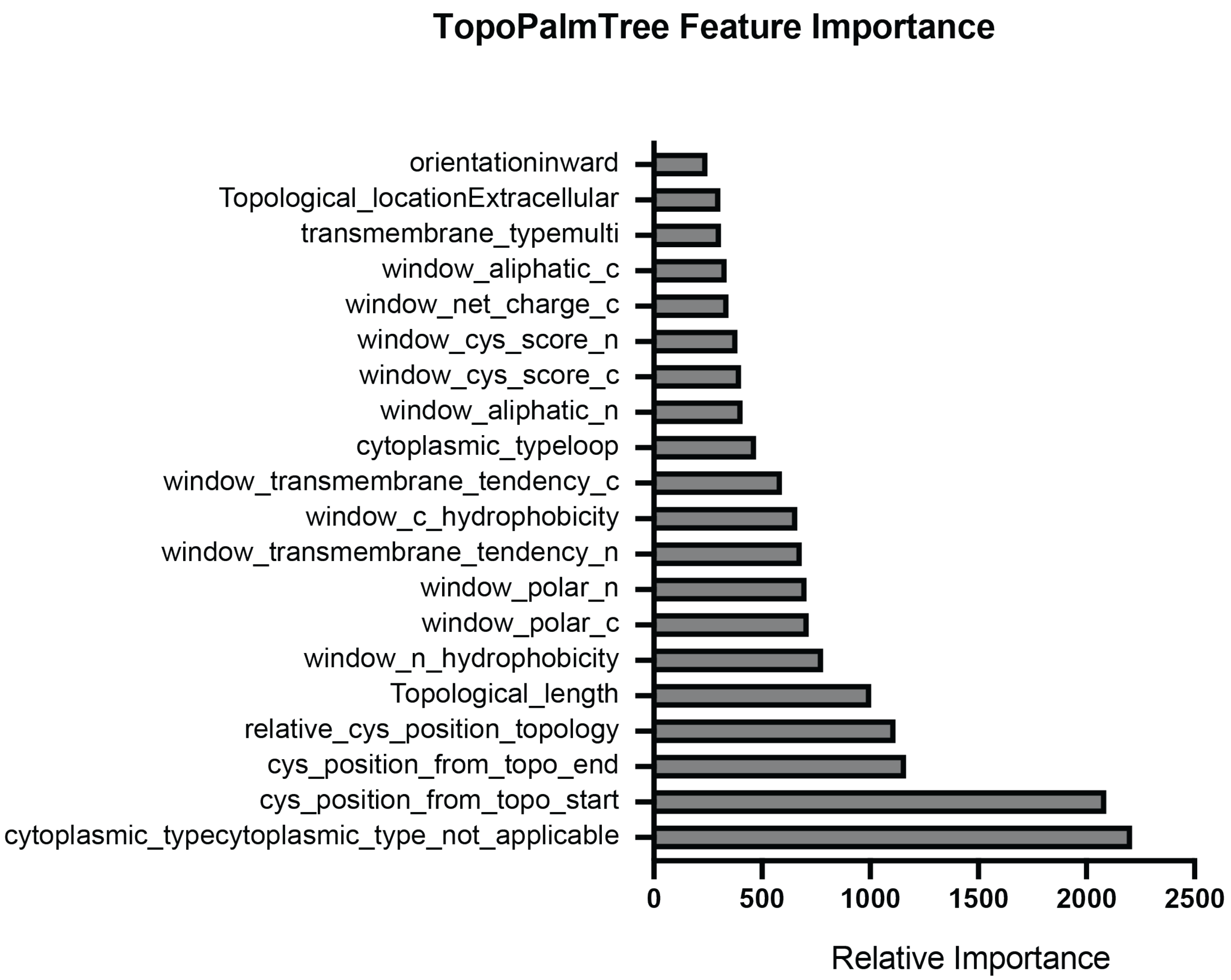

**Figure S5. Feature importance plot for TopoPalmTree.** The varImp function from the caret package was used to examine feature importance for the top 20 features. Importance is determined by reduction in the Gini index. The highest scoring feature determines whether the Cys site is cytoplasmic. Features ending with _n and _c represent the N- and C-terminal windows, respectively, for the indicated feature.

SUPPLEMENTARY TABLES

**Table S1. Results of TopoPalmTree applied to the holdout viral *S*-palmitoyl protein dataset.** Class1 probability reflects the TopoPalmTree output score for a positive class (i.e. *S*-palmitoyl) assignment.

**Table S2. TopoPalmTree applied to the mouse transmembrane proteome.** Class1 probability reflects the TopoPalmTree output score for a positive class (i.e. *S*-palmitoyl) assignment. A total of 49828 Cys residues were amenable to inference by TopoPalmTree. A minority of the Cys residues (5562) were located in rare locations (e.g. signal sequences) not represented in the training dataset and are considered not amenable to inference. Those sites and any other sites not amenable to inference have NA values in the probability column.
